# Consistent movement of viewers’ facial keypoints while watching emotionally evocative videos

**DOI:** 10.1101/2024.05.02.592052

**Authors:** Shivansh Chandra Tripathi, Rahul Garg

## Abstract

Neuropsychological research aims to unravel how diverse individuals’ brains exhibit similar functionality when exposed to the same stimuli. The evocation of consistent responses when different subjects watch the same emotionally evocative stimulus has been observed through modalities like fMRI, EEG, physiological signals and facial expressions. We refer to the quantification of these shared consistent signals across subjects at each time instant across the temporal dimension as Consistent Response Measurement (CRM). CRM is widely explored through fMRI, occasionally with EEG, physiological signals and facial expressions using metrics like Inter-Subject Correlation (ISC). However, fMRI tools are expensive and constrained, while EEG and physiological signals are prone to facial artifacts and environmental conditions (such as temperature, humidity, and health condition of subjects). In this research, facial expression videos are used as a cost-effective and flexible alternative for CRM, minimally affected by external conditions. By employing computer vision-based automated facial keypoint tracking, a new metric similar to ISC, called the *Average t-statistic*, is introduced. Unlike existing facial expression-based methodologies that measure CRM of secondary indicators like inferred emotions, keypoint, and ICA-based features, the *Average t-statistic* is closely associated with the direct measurement of consistent facial muscle movement using the Facial Action Coding System (FACS). This is evidenced in DISFA dataset where the time-series of *Average t-statistic* has a high correlation (*R*^2^ = 0.78) with a metric called *AU consistency*, which directly measures facial muscle movement through FACS coding of video frames. The simplicity of recording facial expressions with the automated *Average t-statistic* expands the applications of CRM such as measuring engagement in online learning, customer interactions, etc., and diagnosing outliers in healthcare conditions like stroke, autism, depression, etc. To promote further research, we have made the code repository publicly available.

## Introduction

A fundamental inquiry in the field of neuroscience revolves around how the brains of different individuals function in a similar manner.^1^ To illustrate this, Hasson et al. [1] conducted a study wherein different subjects watched a popular movie while undergoing brain imaging. Their research uncovered significant synchronization of brain activity among these individuals when they were exposed to emotionally engaging scenes. Hasson et al. [2] demonstrated that certain films possess the ability to exert substantial control over both brain activity and eye movements, leading to synchronized temporal responses among viewers. Similar consistent response has been observed in other modalities such as EEG [3, 4], and physiological signals such as HRV/ECG [5–7], skin conduction [6–9] when different subjects are exposed to the same emotionally evocative stimulus. We refer to the quantification of this shared consistent response (fMRI/EEG, physiological indicators, facial expressions, etc.) across different subjects exposed to the same (emotionally evocative) stimulus as Consistent Response Measurement (CRM).

CRM is widely explored with fMRI neuroimaging data. Hasson et al. [2] introduced a novel method for evaluating the influence of films on viewers’ brain activity using functional magnetic resonance imaging (fMRI) and inter-subject correlation analysis (ISC). ISC offers a quantitative approach for assessing the degree of similarity in brain activations across individuals. Hasson et al. [10] further applied ISC analysis to autism and found that the stimuli evoke highly shared responses in typical individuals (high ISC), while the response was more variable across individuals with autism (low ISC). ISC analysis has also been explored to differentiate ADHD [11] and depressed subjects [12] from healthy individuals. fMRI-based ISC analysis has been applied to numerous other studies, for studying different brain dynamics, different stimulating environments and so forth [13–22]. Some other metrics based on fMRI includes Inter-subject Functional Correlation (ISFC) [23] and Inter Subject Temporal Synchronization Analysis (IS-TSA) [24] to study CRM in the brain’s Default Mode Network (DMN). While ISC focuses on intra-regional correlations across brains exposed to the same stimulus, ISFC also considers inter-regional correlations. IS-TSA reveals lower DMN synchronization compared to early sensory task-positive regions during attention-demanding stimuli.

CRM has also been explored for EEG data. Dmochowski et al. [3] utilized EEG data with ISC analysis and found that correlated components of EEG occur with emotionally arousing moments of the films. Further, Dmochowski et al. [25] noted that inter-subject correlation (ISC) in EEG responses possesses predictive capabilities, enabling the identification of stimuli that individuals find favorable. Additionally, other studies [4, 26] leveraged EEG data and proposed CRM metrics, such as the “impression index” to assess the impressiveness of a video scene, and the “global synchronization index” to monitor the emotional arousal of a population during video watching. Some of the studies have also utilized physiological signals such as Heart Rate Variability (HRV), Galvanic Skin Response (GSR), ECG (Electrocardiogram), and Cardiovascular signals for CRM analysis using ISC metric [5–9]. Consequently, these studies indicate that CRM opens up interesting applications such as finding evocative parts of a movie/film and the impact of film on viewers’ mind [2, 4], identifying an outlier from typical individuals such as autism [10], depression [12], ADHD [11], Alzheimer’s, Parkinson’s, and Schizophrenia etc. [27].

Although neuroimaging tools like fMRI are popular for CRM studies, they have drawbacks [4]. First, fMRI setups differ from real-life settings as participants are confined in scanners, which might not evoke genuine emotions. Second, fMRI accommodates only one viewer, while film experiences can vary in group settings. Third, fMRI is costly and lacks portability, making it less preferred compared to EEG and physiological methods [28–30]. EEG is often employed as a lower-cost method, offering better temporal resolution (at the cost of limited spatial resolution), and can be carried out in an open environment. However, contamination of EEG signals by extra-cerebral artifacts, facial muscle movements, and eyeblink is a well-recognized problem [31–38]. Further, other physiological signals such as HRV, GSR, ECG, etc. offer greater flexibility for CRM compared to neuroimaging techniques, however, they are susceptible to various factors, including environmental conditions such as temperature and humidity and the health conditions of the subjects [39].

On the other hand, facial expression-based CRM are more flexible since recording facial expression videos only requires a high-resolution camera. In contrast to fMRI, it provides an open recording environment for single or multiple viewers, remains unaffected by external artifacts like eye or facial muscle movements, unlike EEG, and is less affected by environmental conditions than physiological signals. Several studies have leveraged facial expressions to measure CRM. For instance, Mauss et al. [8] recorded facial expressions and physiological signals when different subjects watched an emotional video. They utilized expressions to estimate perceived and experienced amusement/sadness intensities and explored temporal correlations of these intensities with physiological signals. Mangus et al. [40] utilized face-tracking based features (e.g. inferred emotions) and fMRI data of the same subjects, revealing high ISC in face-tracking features for disposition-inconsistent stimuli and in fMRI signals for disposition-consistent stimuli. Other studies also utilized facial expression-based CRM and the ISC metric to investigate differences in temporal synchronization of facial features during solo versus dyad video-watching [41], the ability of facial thermal images to capture emotion-related changes induced by stimuli [42], and whether the synchrony of inferred emotions predicts temporal engagement of audience during theatrical performances [43].

However, the facial expression method in Mauss et al. [8] relied on manually coded emotion ratings which is labor-intensive. Additionally, in the studies [40– 43], automated methods primarily focused on evaluating the consistency of secondary indicators. These indicators included deep learning/commercial software-based estimated emotions, facial keypoint-based distance features, or facial thermal images-based ICA components. The emphasis was on measuring the consistency of these indicators rather than determining whether the facial expressions were actually elicited consistently across subjects. The use of CRM, which can indicate whether expressions were consistently elicited, can offer a more reliable insight into the synchronization of subjects in video watching. Just as synchronized neuronal responses can indicate the viewers’ level of engagement, with deviations suggesting lower engagement [3], we hypothesize that consistent facial expressions can be a viable metric for quantifying engagement. Moreover, given the ease of recording facial expressions, it may efficiently measure engagement across applications such as online learning for students, customer interactions in business, movie watching, patient engagement in healthcare, and so forth.

The present study advances and refines prior methodologies for assessing CRM through facial expressions. It introduces a new metric, termed the *Average t-statistic* which is automated and closely associated with measuring the consistency of facial expressions utilizing the Facial Action Coding System (FACS) [44, 45]. This metric relies on the statistical modeling of facial keypoints, which can be automatically tracked using computer vision or deep learning models. To validate its effectiveness in indicating the consistency of expressions, we introduce a metric called *AU consistency* which involves manual FACS coding of each video frame to quantify CRM of facial muscle movements at each time point. Subsequently, we demonstrate that although the *Average t-statistic* is not directly reliant on FACS coding, it exhibits a robust *R*^2^ value of 0.78 with *AU consistency* in the DISFA dataset.

The subsequent section provides a detailed definition of the metrics that utilize facial expression videos to quantify CRM. Further, we discuss the experimentation and evaluation on the DISFA dataset, and finally, we summarize our work’s limitations and outline potential future directions.

## CRM using facial expressions

To explore the consistent elicitation of facial expressions, we employ the publicly accessible DISFA dataset [46, 47]. This dataset consists of 27 subjects who viewed the same emotionally evocative video lasting approximately 4 minutes and 1 second, during which their facial expressions were recorded at a rate of 20 frames per second (fps). Our initial step involves the establishment of the *AU consistency* metric, designed to gauge CRM by leveraging FACS coding for 12 Action Units (AUs) at each frame of the subjects in the DISFA dataset (Table 1). Following that, we introduce our keypoint-based metrics and statistical models to quantify CRM at each time point. The following sections provide a detailed explanation of the proposed facial expression-based metrics for CRM.

**Table 1.** Description of the 12 AUs coded in DISFA

| AU | Name | Appearance Change |
| --- | --- | --- |
| 1 | Inner Brow Raiser | InnerBrows are pulled upwards |
| 2 | Outer Brow Raiser | OuterBrows are pulled upwards |
| 4 | Brow Lowerer | Both brows are pulled downwards- sometimes only the InnerBrows, or including the central or Outer-Brows |
| 5 | Upper Lid Raiser | Upper eyelid is raised |
| 6 | Cheek Raiser | Skin from cheeks and temples are pulled towards the eyes |
| 9 | Nose Wrinkler | Moves skin around nose towards nasal root causing wrinkles |
| 12 | Lip Corner Puller | Lifts the lip corners obliquely upwards |
| 15 | Lip Corner Depressor | Pulls lip corner down |
| 17 | Chin Raiser | Pushes the lower lip and chin upward |
| 20 | Lip Stretcher | Pulls lip corners horizontally outwards |
| 25 | Lips Part | Upper and lower lip are separated |
| 26 | Jaw Drop | Jaw moves down |

**Table 2.** Start and end frame number of different target emotion segments

| Start Frame No. | End Frame No. | Target Emotion |
| --- | --- | --- |
| 65 | 283 | Happy1 |
| 343 | 755 | Happy2 |
| 817 | 1,165 | Surprise1 |
| 1,227 | 1,354 | Fear |
| 1,414 | 1,874 | Disgust1 |
| 1,935 | 2,455 | Disgust2 |
| 2,515 | 3,200 | Sadness1 |
| 3,261 | 3,934 | Sadness2 |
| 3,995 | 4,832 | Surprise2 |

### CRM using FACS coding

Fine-grained movement of facial expressions can be directly labeled through visual inspection by trained experts. One of the most widely accepted systems for such measurement is the Facial Action Coding System (FACS) [45] developed by Ekman and Friesen. FACS is a comprehensive facial coding system to taxonomize different facial muscle movements based on their appearance. The taxonomized movements are called Action Units (AUs). An Action Unit (AU) constitutes the activation of a single facial muscle or a specific group of muscles that always move together as a unit. Any facial expression can be encoded as a combination of a single or a group of AUs. Since AUs give a direct measurement of facial movements without any secondary resort in between, we use AUs directly to efficiently find consistent expressions. We define our *AU consistency* metric that utilizes the FACS coding as follows:

### *AU consistency* metric (*κ*_*AU*_)

Consider facial video recordings of *n* subjects watching the same emotionally evocative video. The *AU consistency* metric utilizes the AU labeling of all the frames of the facial video recordings of the subjects. Let *μ*(*i, k, τ*) represent the AU labeling for the *k*th AU in the facial video of *i*th subject at time instant *τ*. *μ*(*i, k, τ*) = 1 if facial video of subject *i* has AU *k* present at time *τ*. The *AU consistency* for AU *k* at time *τ* is defined as 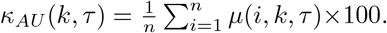.The overall *AU consistency* at time *τ* is defined as *κ*_*AU*_ (*τ*) = max *κ*_*AU*_ (*k, τ*).

In summary, a higher *AU consistency* indicates that a significant number of subjects exhibit the same AU and therefore the same facial muscle movement at a given time point, whereas a lower value indicates fewer subjects displaying the same facial muscle movement. Note that *AU consistency* measures the degree of consistent facial expressions rather than quantifying the number of facial components exhibiting consistency. For instance, if *AU consistency* reaches 100 percent, it signifies one AU is consistently present across all subjects, regardless of whether it involves just one eyebrow or the entire face. In both cases, the degree of consistency remains the same.

### CRM using facial keypoint tracking

For an automated facial expressions analysis in a video sequence, we identify and track specific keypoints located at various landmarks on the face, including the eyes, eyebrows, nose, lips, and jawline (Fig. 1). The labeling of these keypoints can be done either manually or through computer vision-based algorithms [48]. We rely on the facial keypoints in the publicly available DISFA dataset [46, 47], which consists of 66 keypoints (Fig. 1) at each of the 4845 video frames of any subject. These keypoints are tracked using the Active Appearance Model (AAM) [49]. We observed that these 66 keypoints are reasonably stable, and accurate, making them suitable to our analysis. Recent advancements in keypoint-tracking technology, incorporate additional facial features such as edge contours [50] or optical flow from video data [51], which may lead to more stable tracking. Once these techniques are incorporated in higher resolution tracking algorithms [52, 53] and give stable and consistent tracking, the number of keypoints may be expanded to 106 or more, thereby enabling the capture of more intricate facial expressions and features, including wrinkles, cheekbone height, and dimples.

**Fig. 1.**
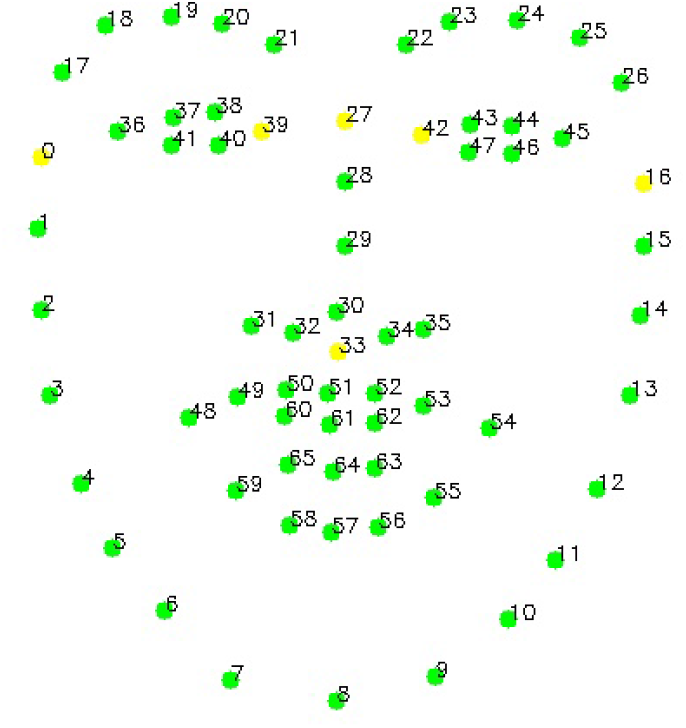
Demonstrating 66 facial keypoints on an in-house subject, akin to those tracked on DISFA subjects. Out of those, six keypoints (yellow-colored) are used for registration using affine transformation.

To ensure uniformity across different subjects and mitigate geometric variations caused by factors such as head motion, variations in face sizes, and relative facial part positions, we preprocess these keypoints [54]. Subsequently, we convert these keypoints into Keypoint Movement (KPM) vectors, representing the movement of keypoints from a neutral frame. Additionally, we establish metrics using statistical modeling on the KPM vectors to quantify CRM as the degree of consistent keypoint movement at each time point. We will first provide a detailed explanation of the geometric corrections:

### Geometric corrections

The process of eliminating variations in facial geometry involves three crucial steps: first, frontalization is carried out, and then registration is performed using affine and similarity transformations, as detailed below:

To isolate facial muscle movements and eliminate the impact of head movement, we employ the algorithm proposed by Vonikakis et al. [55]. This algorithm transforms facial keypoints to resemble a front-facing image.

Following this, an affine registration process [56] is employed to align the keypoints with a standard reference, reducing variations related to face size and position. We specifically choose six fixed facial keypoints to estimate geometric parameters within a facial image. These parameters are then used to register the remaining keypoints in the image using an affine transformation with six parameters. In the case of DISFA, the selected six keypoints are: 0, 16, 39, 42, 27, and 33 (Fig. 1).

Lastly, we apply similarity registration [56] to address intra-face variations, such as the distance between eyebrow corners, nose length, eye dimensions, and more. We perform this registration face part-wise using specific fixed keypoints. For instance, keypoints 42 and 45 are used for the left eyebrow and left eye, 36 and 39 for the right eyebrow and right eye, 27 for the nose, and, 0 and 16 for the jawline. Each of these similarity transformations involves four parameters. Lips, however, lack fixed keypoints and are consequently not subjected to similarity registration.

### Keypoint Movement (KPM) vector

A Keypoint Movement (KPM) is a vector that characterizes motion in facial keypoint positions relative to a neutral face. To generate a KPM vector, we compute the difference between the x and y coordinates of the preprocessed facial keypoints across any frame with a neutral frame of the same subject. The KPM of a subject at time *τ* is represented as a vector *X*(*τ*) *∈* ℝ ^132^.

### Statistical modeling of KPM

At each timepoint *τ*, the KPM vector *X*(*τ*) is assumed to be a multivariate random variable. The KPMs of different subjects are considered to be independent and identically distributed (iid) samples from the same underlying time-dependent distribution. We perform hypothesis testing under different statistical models to evaluate whether the facial keypoints exhibit significant movement from the neutral position. The null hypothesis being tested is that the mean KPM vector at *τ* is equal to the zero vector (a neutral face keypoint position). The results of these hypothesis tests are then transformed into *t*-scores. At any given *τ*, these *t*-scores are further mapped to a specific function definition (Average, Maximum, etc.), which serves as a single metric value for quantifying the degree of consistent keypoint movements. Like *AU consistency*, these metrics measure the degree of consistent keypoint movements, rather than quantify how many keypoints are consistent, i.e., a high metric value implies that keypoints are consistent (irrespective of whether few are consistent or all are consistent). In the following sections, we provide a detailed explanation of each of the statistical models and the metrics proposed under them:-

1. **Independent Univariate Gaussian Model -** In the independent univariate Gaussian model, we assume that each random variable *X*_*i*_(*τ*) (where *i* ranges from 1 to 132) in the KPM vector *X*(*τ*) follows a normal distribution with a mean of *μ*_*i*_(*τ*) and a variance of 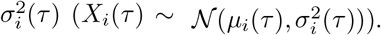. Under the null hypothesis *H*_0_(*τ*)^*i*^ for component *i* there is no discernable consistent movement of the keypoints in any direction *H*_0_(*τ*)^*i*^ : *μ*_*i*_(*τ*) = 0. To test the null hypothesis, we employ a two-tailed Student’s t-test. The t-score for each component *i* is calculated as 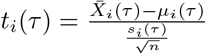 (where 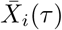 is sample mean and *s_i_* (*τ*) is sample standard deviation). This process results in a 132-dimensional *T* (*τ*) vector of t-statistic values for a given timepoint *τ*. The magnitude of these t-scores indicates the significance of the hypothesis test, while the sign only indicates the direction of movement, i.e., whether the corresponding x or y coordinate of the keypoint moves in a positive or negative direction. We are primarily interested in the magnitude of the t-scores to gauge the strength of consistent keypoint movements within a population. We propose two metrics based on the magnitude of the t-scores:
  a. ***Average t-statistic***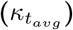: The *average t-statistic* is calculated as the mean of the magnitudes of t-scores of all the keypoints at a specific timepoint *τ*. Formally, 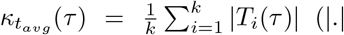 denotes absolute value). This metric gives a good indication of CRM if a large number of keypoints are consistent.
  b. ***Maximum t-statistic***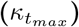 : The *Maximum t-statistic* is defined as the maximum magnitude among all the keypoint t-scores. Formally, 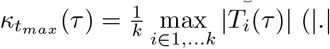 denotes absolute value). In situations where only a few keypoints exhibit consistent movement (e.g., only eyebrows), and others do not, the average t-statistic value may be extremely low, indicating wrongly that there are no consistent keypoint movements. Therefore, the *Maximum t-statistic* will capture the degree of consistency in this case by picking up only the maximum value.
2. **Independent Bivariate Model -** In the independent bivariate model, the movement along the *x* and *y* directions of any keypoint is considered as a bi-variate normal distribution. Let *r*_*i*_(*τ*) = (*X*_2*i−*1_(*τ*), *X*_2*i*_(*τ*)) represent the movement vector of the *i*th keypoint at time *τ* with respect to the neutral face. In this model, *r*_*i*_(*τ*) *∼* 𝒩 (*μ*_*i*_(*τ*), Σ_*i*_(*τ*)), where *μ*_*i*_(*τ*) *∈* ℝ ^2^ is a vector of means and Σ_*i*_(*τ*) *∈* ℝ ^2*×*2^ is a covariance matrix. The null hypothesis is given by *H*_0_(*τ*)^*i*^ : *μ*_*i*_(*τ*) = (0, 0), which examines whether the keypoints exhibit any significant movement from their neutral position. To evaluate this hypothesis, we employ a Hotelling T-square test, which is a generalization of the Student’s t-test to multivariate hypothesis testing [57]. This test calculates a Hotelling *t*^2^-score as follows: 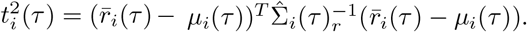. Here, 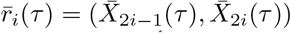 represents a vector containing the sample means, 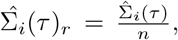, and 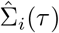 is the sample covariance matrix. By applying this test to all 66 keypoints at a specific timepoint *t*, we obtain 66-dimensional vector *T* ^2^(*τ*) of *t*^2^-scores, each reflecting the significance of movement for the corresponding keypoint. To quantify CRM under this model using consistent keypoint movements, we propose two metrics that are similar to metrics proposed under the independent univariate model:
  a. ***Average*** *t^2^* ***-statistic***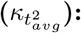 : The *average t*^2^*-statistic* is calculated as the mean of all *t*^2^-statistics at timepoint *t*. Formally, 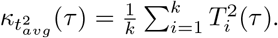
  b. ***Maximum*** *t^2^* ***-statistic***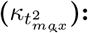 : The *Maximum t*^2^*-statistic* represents the highest among all the *t*^2^-statistics at timepoint *t*. Formally,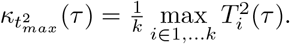. In scenarios where only a few keypoints exhibit consistent movement, the maximum *t*^2^-statistic makes a better assessment than the average *t*^2^-statistic.
3. **Multivariate Gaussian with Dimensionality Reduction** (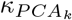) **-** In this model, we assume that the random variables *X*(*τ*) follow a multivariate gaussian distribution, represented as *X*(*τ*) *∼* 𝒩 (*μ*(*τ*), Σ(*τ*)), where *μ*(*τ*) *∈* ℝ ^132^ and Σ(*τ*) *∈* ℝ ^132*×*132^. To test the null hypothesis *H*_0_(*τ*) : *μ*(*τ*) = (0, 0, …..132 times), the Hotelling T-square test may be employed to compute the *t*^2^-statistic using a similar formulation as defined in the independent bivariate model. This single *t*^2^-statistic represents the CRM metric (*κ*_*MV* *G*_(*τ*)) under this model at time *τ* . However, the computation of Hotelling’s *t*^2^-score requires that the number of samples (subjects) significantly exceeds the number of dimensions in the vector *X*(*τ*), or else 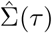 becomes a singular matrix. Therefore, in the case of data from fewer subjects, *κ*_*MV* *G*_(*τ*) may not be computable. To address this dimensionality issue, we apply Principal Component Analysis (PCA) to reduce the dimensions in the given dataset. Let the complete dataset be denoted as *Y ∈* ℝ ^132*×m*^, where *m* is the total number of samples. The PCA decomposition yields an approximation, *Y ≈ PQ*, where *P ∈* ℝ ^132*×k*^ and *Q ∈* ℝ ^*k×m*^ (*k <<* 132). Now, the low-dimensional matrix *Q ∈* ℝ ^*k×m*^ represents the data, and the random variable at time *τ* can be represented as a vector *Q*(*τ*) *∈* ℝ ^*k*^. Assuming, *Q*(*τ*) follows a multivariate Gaussian distribution represented as *Q*(*τ*) *∼* 𝒩 (*μ*(*τ*), Σ(*τ*)) where *μ*(*τ*) *∈* ℝ ^*k*^ and Σ *∈* ℝ ^*k×k*^, we test the null hypothesis *H*_0_(*τ*) : *μ*(*τ*) = (0, 0, …*k times*) using a Hotelling T-square test [57]. The Hotelling *t*^2^-statistic computed under *H*_0_(*τ*) represents the CRM metric at time *τ* known as *t*^2^*-statistic with PCA* or 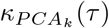 (*k* is the number of PCA components). In the DISFA dataset, we retained the top five components (*k* = 5), which can explain 90 percent of the variance and compute the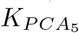.

## Experiments

In this section, we illustrate the application of *AU consistency* (*κ*_*AU*_) using the DISFA dataset to evaluate CRM, by assessing the consistency of expressions in the temporal dimension. We demonstrate how *AU consistency* facilitates the comparison of consistency levels across various emotion segments in the DISFA stimulus. Additionally, considering that *AU consistency* involves manual FACS coding, we engage in a discussion about potential automated keypoint-based metrics that scould serve as viable alternatives.

### DISFA stimulus

The stimulus is 4m 1s in length and consists of nine segments targetting five emotions-Happy, Surprise, Fear, Disgust, and Sadness. If multiple segments target the same emotion, we index them starting from one, e.g., Happy1 and Happy2 (Table. 2). The segments are separated by 2-3 seconds and we call this Inter-Segment Gap (ISG).

### CRM analysis using *AU consistency*

We examine the *AU consistency* in DISFA across the video timeline, as illustrated in Figure 2. This analysis provides insight into the consistency of elicited facial expressions at each of the 4845 time points. Notably, segments like Happy1 and Happy2 demonstrate a high level of *AU consistency* (25.93-96.30%), while Sadness1 and Sadness2 result in expressions with low levels of *AU consistency* (11.11-37.04%). In the case of Happy1, the *AU consistency* gradually builds and peaks towards the end of the segment. Additionally, the high *AU consistency* persists in the Inter-Segment Gap (ISG), suggesting a significant delay before the facial expression returns to a neutral state. This may be verified for subjects in DISFA, see the illustration in Figure 3. The figure reveals that the subject displays facial expressions even in the ISG, and it doesn’t settle into a completely neutral expression. Consequently, we extend the boundaries of the emotion segment into their subsequent ISG for further analysis.

**Fig. 2.**
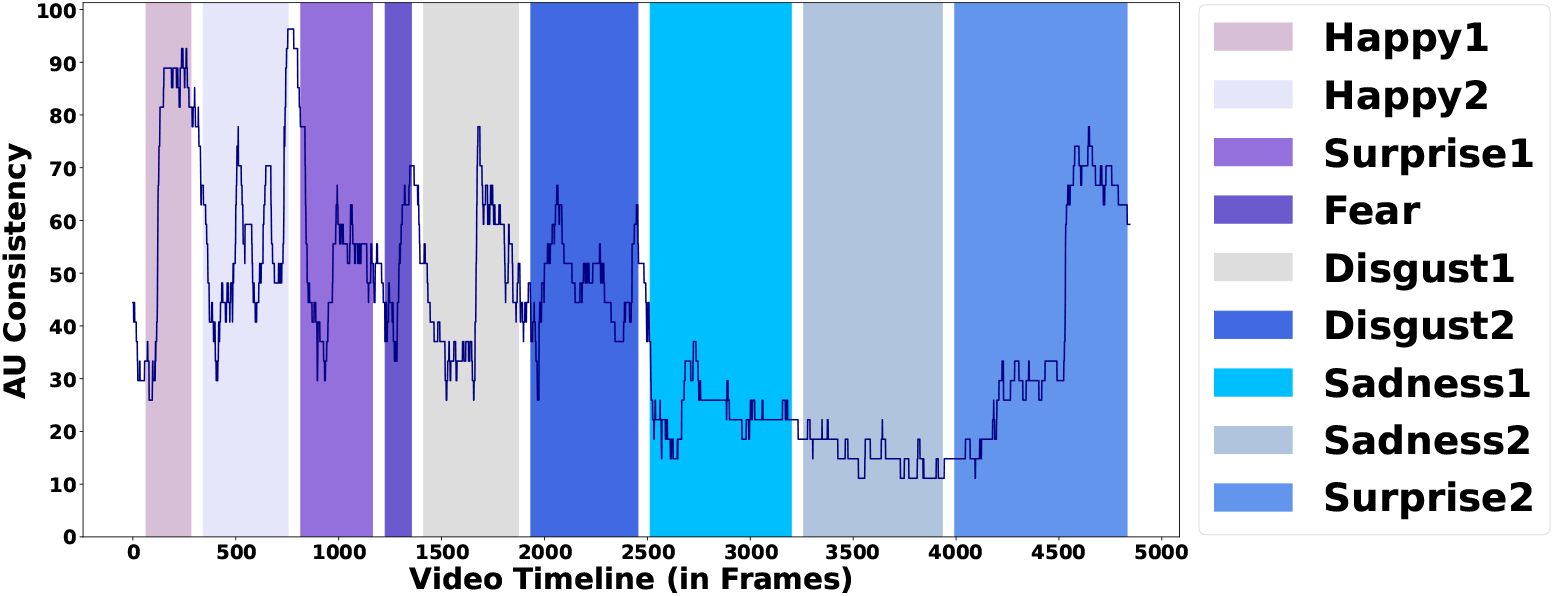
*AU consistency* along the video timeline in DISFA. Colored bars represent the different emotion segments and the interval represents the Inter-Segment Gap (ISG)

**Fig. 3.**
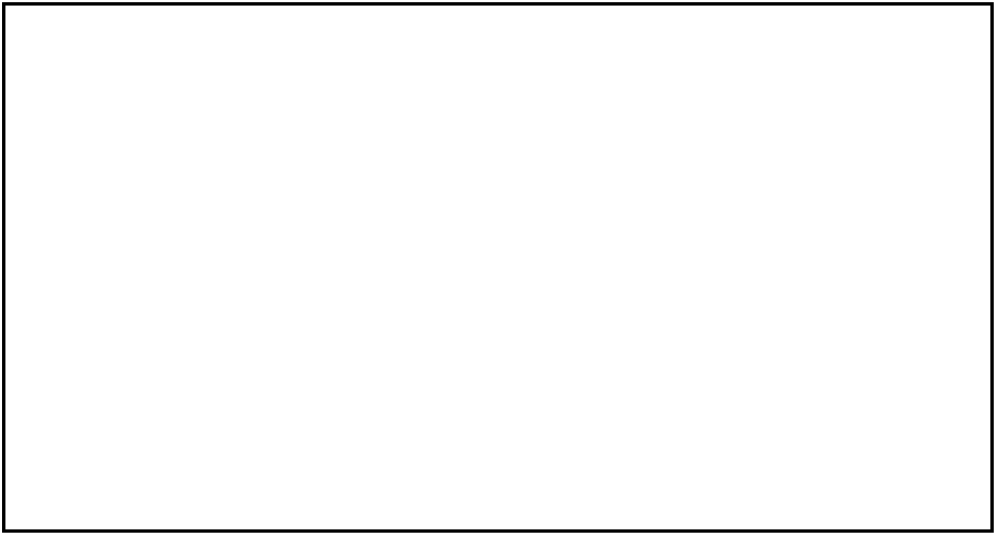
Expression of a subject at the start, peak consistent frame, and end of an emotion segment when watching the DISFA stimulus. This subject was not a part of DISFA dataset and included solely for illustrative purposes, highlighting the non-neutral expressions observed within the Inter-Segment Gap (ISG) of some DISFA subjects.

To facilitate the comparison of various emotion segments, we’ve categorized the 4845 time points into four groups (*Inconsistent, Low Consistent, Mild Consistent, High Consistent*) through an analysis of the Cumulative Distribution Function (CDF) of the *AU consistency* ‘s values (refer to Fig. 4). The CDF exhibits an approximately linear trend for *AU consistency* in the range of [0-70.37%) and a distinct linear trend for *AU consistency* in the range of [70.37-100%]. The latter represents approximately 10 percent of the time points, which we classify as the *High Consistent* category. The remaining graph is evenly divided into three segments, assigned to *Inconsistent, Low Consistent*, and *Mild Consistent* categories, respectively. Refer to Table 3 for the *AU consistency* range corresponding to each class.

**Table 3.** Consistency classes based on *AU consistency*

| AU consistency range(%) | Sample percentile range(%) | Class |
| --- | --- | --- |
| [0,25.93) | [0,30) | Inconsistent |
| [25.93,44.44) | [30,60) | Low Consistent |
| [44.44,70.37) | [60,90) | Mild Consistent |
| [70.37,100] | [90,100] | High Consistent |

**Fig. 4.**
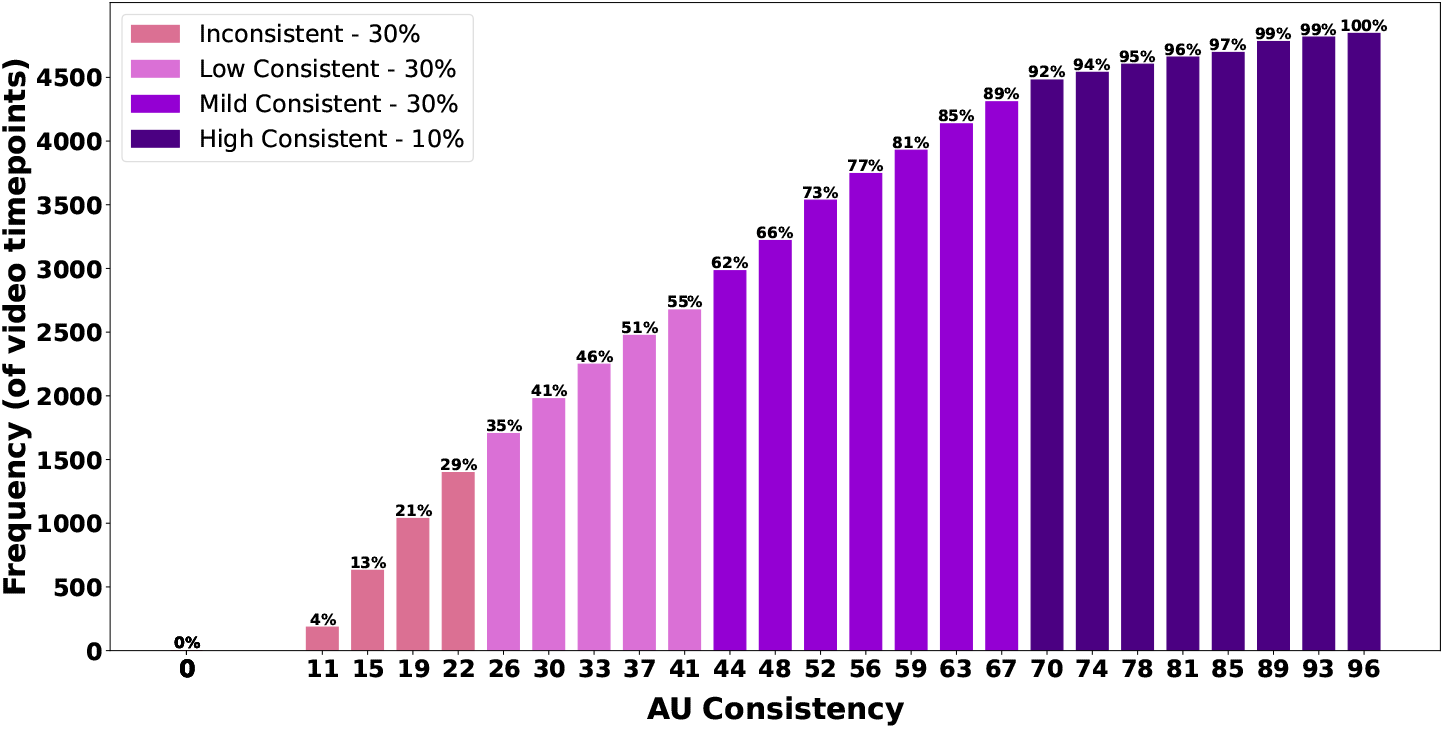
Cumulative Distribution Function (CDF) of the *AU consistency*. For any given bar, the percentage above indicates the proportion of time points (t=1,2,…4845) where the *AU consistency* is less than or equal to the value associated with that bar on the x-axis.

Table 4 reveals the distribution of the four consistency classes within each emotion segment. In Happy1, for instance, 74 percent of frames belong to the *High Consistent* class, the highest percent across all segments. Similarly, Surprise1 and Disgust2 has the highest percent (77) of frames in the *Mild Consistent* class. Disgust1, on the other hand, exhibits the highest percent (45) of frames in the *Low Consistent* class, while Sadness2 exhibits the highest percent (100) of frames in the *Inconsistent* class.

**Table 4.** Distribution of the four consistency classes present in different emotion segments

| Emotion | Inconsistent | Low Consistent | Mild Consistent | High Consistent |
| --- | --- | --- | --- | --- |
| Happy1 | 6 | 14 | 6 | 74 |
| Happy2 | 0 | 14 | 60 | 27 |
| Surprise1 | 0 | 17 | 77 | 6 |
| Fear | 0 | 20 | 69 | 11 |
| Disgust1 | 1 | 45 | 49 | 5 |
| Disgust2 | 1 | 22 | 77 | 0 |
| Sadness1 | 89 | 11 | 0 | 0 |
| Sadness2 | 100 | 0 | 0 | 0 |
| Surprise2 | 33 | 31 | 19 | 16 |

**Table 5.** KL-divergence (row-wise averaged) between *κ*_*AU*_ distribution table and each of the five keypoint-based metrics 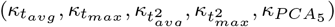

| Keypoint-based metric | Avg. KL-divergence (per row) |
| --- | --- |
| *Average t-statistic* ($\kappa_{t_{avg}}$) | 0.92 |
| *Maximum t-statistic* ($\kappa_{t_{max}}$) | 0.98 |
| *Average t<sup>2</sup>-statistic* ($\kappa_{t_{avg}}^2$) | 0.93 |
| *Maximum t<sup>2</sup>-statistic* ($\kappa_{t_{max}}^2$) | 0.98 |
| *t<sup>2</sup>-statistic with PCA* ($\kappa_{PCA_5}$) | 0.96 |

These findings suggest that not all emotions elicit consistent facial expressions to the same extent, and the Happy emotion may elicit the highest degree of consistency. However, we observe that within emotion, the amount of consistency varies for different segments. For instance, Happy1 and Happy2 differ significantly in the *High Consistent* class. This may be possible due to the stimulus choice and design within a segment.

The analysis above demonstrates that *AU consistency* can be a suitable metric for gauging the level of CRM during the viewing of emotionally evocative videos. Furthermore, it offers a means to evaluate the effectiveness of various emotion segments within a stimulus video. This metric relies on FACS coding, a method that involves the precise quantification of subtle facial muscle movements by certified FACS coders. Consequently, we hypothesize that *AU consistency* stands as a highly reliable metric for quantifying CRM using facial expression videos.

### Measuring CRM using facial keypoints

FACS coding, while effective, can be costly and time-consuming, making it impractical in real-time situations. A more feasible approach involves creating metrics that rely on the automated tracking of facial expressions using facial keypoints based on computer vision or deep learning algorithms. This renders an automated system that offers the advantage of real-time measurements of consistent response.

To determine the most reliable alternative to the *AU consistency* metric among the proposed keypoint-based metrics, we conducted a linear regression analysis. This analysis examines the relationship between the *AU consistency* metric and five keypoint-based metrics across the video timeline. The resulting best-fit line is illustrated in red in Fig. 5. Coefficient of determination *R*^2^ is computed, to assess the goodness of fit for each best-fit line.

**Fig. 5.**
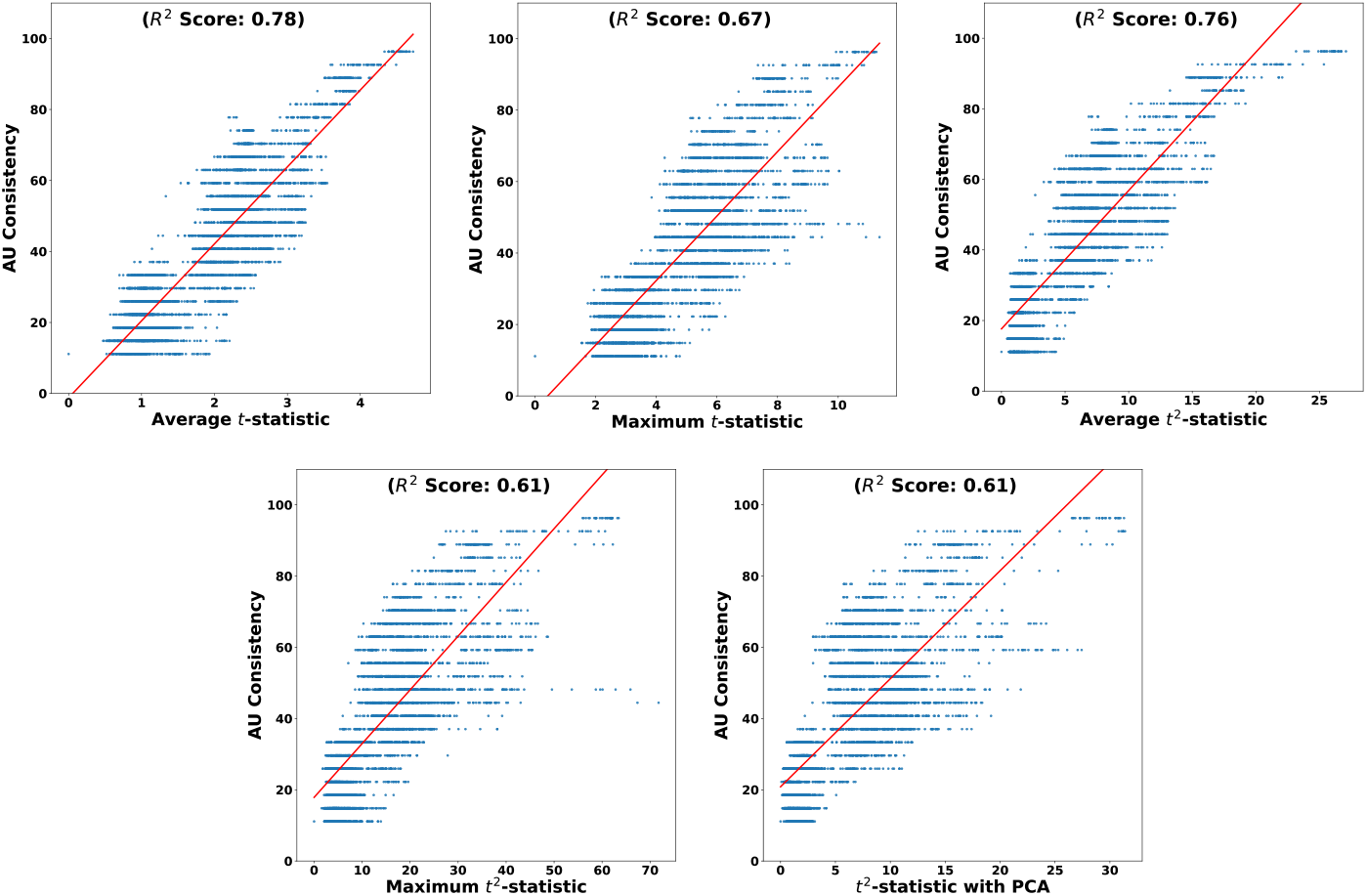
Regression lines depicting the correlation between *AU consistency* and different keypoint-based metrics, using their 4845 data points across the video timeline.

The maximum *R*^2^(=0.78) is obtained by the metric *Average t-statistic*. Figure 6 further demonstrates the similarity in trends between the *AU consistency* metric and the *Average t-statistic* metric along the video timeline in the DISFA dataset.

**Fig. 6.**
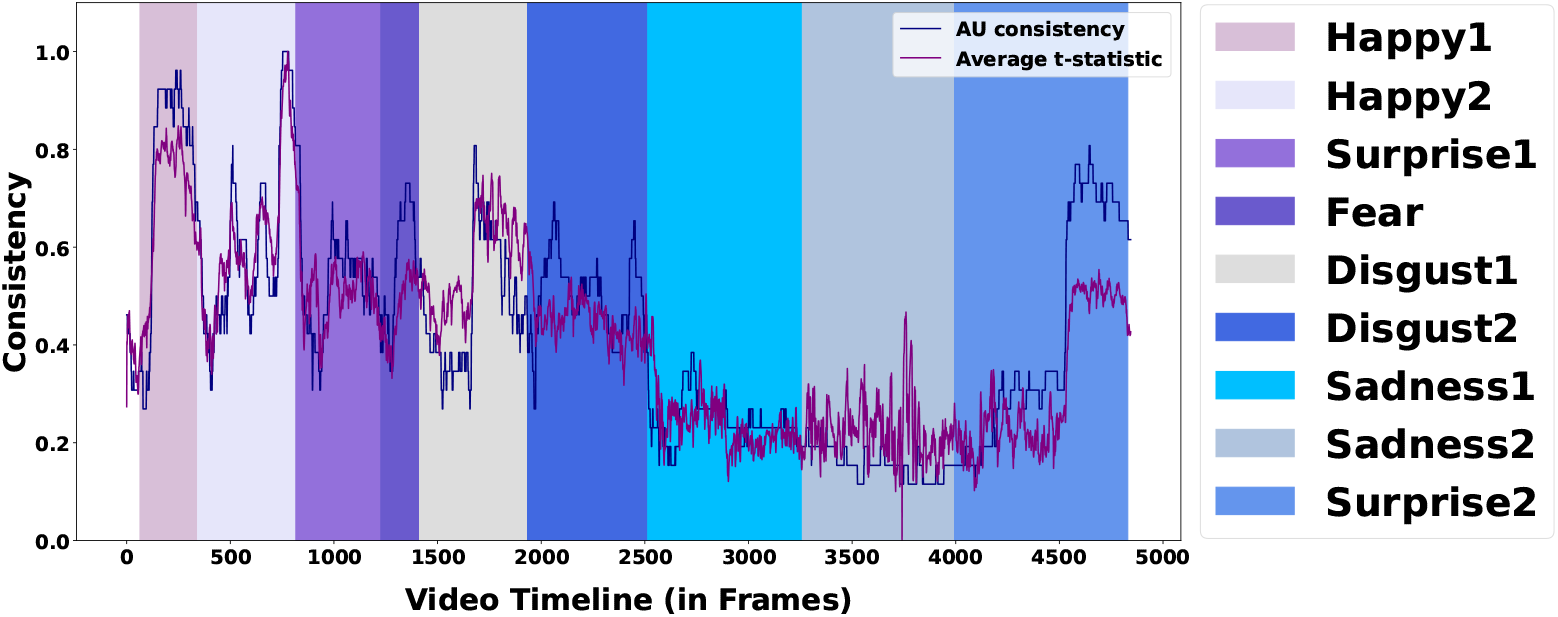
*AU consistency* and *Average t-statistic* along the video timeline (y values normalized for both metrics between [0,1]). Colored bars represent the emotion segments extended to their subsequent ISGs.

### Consistency class distribution

We investigated whether the distribution of keypoint-based metrics across four consistency classes (*High Consistent, Mild Consistent, Low Consistent*, and *Inconsistent*) aligns with the distribution of *AU consistency*. To achieve this, we generated a table for each keypoint-based metric similar to the structure of Table. 4 created for the *AU consistency* metric. We first identified the range of each keypoint-based metric for specific sample percentile intervals associated with the four classes, as outlined in Table. 3 (ranges: [0,30), [30,60), [60,90), and [90,100]). For instance, let’s consider 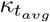. The respective ranges for 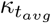 at these percentile intervals are [0,1.19), [1.19,2.22), [2.22,3.09), and [3.09,4.72]. Using these ranges, we constructed a table analogous to Table. 4 for the 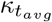 metric. We repeated this process for other keypoint-based metrics and consolidated all the distributions in one table (Table. 6).

For any metric, each row sums up to 100 or dividing the row by 100 transforms it into a probability distribution. Kullback-Leibler (KL) Divergence serves as a measure to quantify the difference between two probability distributions. To identify which keypoint-based metric distribution in the table most closely resembles that of *κ*_*AU*_ distribution, we employed the average Kullback-Leibler (KL) Divergence per row. For instance, we took the Happy1 row for *κ*_*AU*_ distribution and the Happy1 row for 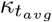 and compute the KL-divergence. This process was repeated for all rows in the 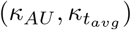 pair, and the average KL-divergence was reported (Table. 5). This procedure was iterated for all keypoint-based metrics, and results illustrate that 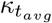 exhibited the least average KL-divergence, thereby demonstrating the closest resemblance to the *κ*_*AU*_ metric.

In summary, the distribution analysis of the four classes shows that *Average t-statistic* represent the *AU consistency* more closely than the other keypoint-based metrics. Moroever, under the assumptions of a linear regression model, the *Average t-statistic* demonstrated the highest *R*^2^(=0.78) with the *AU consistency* metric. Therefore, we recommend using the *Average t-statistic* as a reliable method for identifying consistent responses based on facial keypoints.

## Conclusion

We demonstrate the effective use of facial expression videos to quantify Consistent Response Measurement (CRM) when multiple individuals watch the same emotionally evocative videos. Unlike existing methods that assess the consistency of secondary indicators such as emotions, keypoint, and ICA-based features, our CRM, based on the *AU consistency* metric utilizes the Action Unit (AU) information based on the FACS to quantify the CRM. The *Average t-statistic* metric, is associated with the direct measurement of consistent facial muscle movements using automated keypoint tracking algorithms. There is a strong correlation (*R*^2^ = 0.78) between the *Average t-statistic* based on keypoint tracking and the *AU consistency* metric that relies on FACS coding of video frames.

Our CRM metrics are cost-effective and more adaptable compared to conventional neuroimaging methods involving expensive fMRI/EEG data. It is also less sensitive than physiological signals such as HRV, GSR, and ECG. This automated *Average t-statistic* metric and its associated statistical model can be applied to quantify observer engagement in videos, identify outlier subjects, and offer insights into various applications. Given the simplicity of recording facial expressions, the metric can be employed to discover emotionally evocative segments in movies, assess the impact of films on viewers’ minds, measure engagement in online learning, evaluate customer interactions in business, assess patient engagement in healthcare, and so forth. Further, it may identify outliers in populations to indicate conditions like stroke, ADHD, Alzheimer’s, Parkinson’s, and Schizophrenia, as well as mental disorders such as autism and depression. Our metric may also assist in video-EEG paradigms, for instance, by quantifying facial expression abnormalities to estimate instants of epileptic seizures and cognitive impairment, such as dementia.

CRM extends its applicability beyond the confines of video-watching stimuli, showcasing its versatility to quantify consistent facial expressions across diverse sensory modalities. For example, it has been utilized to assess synchronized neural responses of subjects when exposed to affective audios and narrated stories [58, 59], affective images [60], olfaction [61], sweet, bitter or salty gustation [62] and so forth. One notable application of CRM may involve quantifying event-related facial expressions (ERFEs), akin to the analysis of event-related potentials (ERPs) in EEG studies [63]. By comparing ERFEs and ERPs, researchers can determine the lag between neural processing and the onset, apex duration, or offset of consistent facial expressions. We hypothesize that variations in these lag can provide valuable indicators of neural processing delays, which may be used to further study and compare characteristics of different population groups, such as healthy, autism, ADHD, or stroke populations. Thus, CRM emerges as a comprehensive tool, capable of quantifying consistent responses across a spectrum of stimuli types and sensory experiences, enriching our understanding of emotional processing and its implications.

Certain factors, such as precise alignment of stimulus presentation and recorded video, are crucial for achieving consistent facial expressions in CRM. Moreover, response time variations to the same stimulus among subjects necessitate careful demographic selection. For example, a homogeneous group, like same-age, same-gender healthy individuals, is preferable to identify evocative segments of a video clip. However, characterizing differences in facial expressions among different population groups, such as normal healthy, ADHD, autism, or stroke patients, may necessitate studying heterogeneous groups of the population. Finally, to enhance the metric’s robustness, we may simultaneously record facial expressions and neuroimaging data, providing insights into both facial responses and direct brain activity while subjects are exposed to a variety of stimuli. The proverb “The face is a picture of the mind” can be literally tested rigorously by correlating the facial keypoint movements with the brain activity obtained using neuroimaging data.

## APPENDIX

Table. 6 shows the distribution of *AU consistency* and the keypont-based metrics in the four consistency classes - *Inconsistent, Low Consistent, Mild Consistent* and *High Consistent* in different emotions.

**Table 6.**
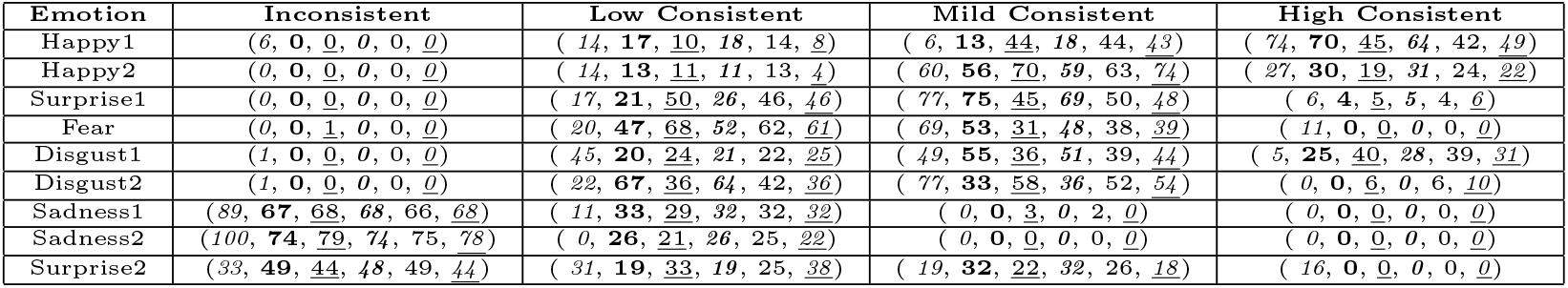
Distribution of the CRM metrics in the four consistency classes per emotion. Each entry contains values in the order 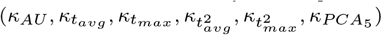

## Footnotes

1 Code available here: https://github.com/Shivansh-ct/Consistent-Keypoints

